# Contextual incongruency triggers memory reinstatement and the disruption of neural stability

**DOI:** 10.1101/2022.07.26.501077

**Authors:** Xiongbo Wu, Pau Packard-Blasco, Josué García-Arch, Nico Bunzeck, Lluís Fuentemilla

## Abstract

Schemas, or internal representation models of the environment, are thought to be central in organising our everyday life behaviour by giving stability and predictiveness to the structure of the world. However, when an element from an unfolding event mismatches the schema-derived expectations, the coherent narrative is interrupted and an update to the current event model representation is required. Here, we asked whether the perceived incongruence of an item from an unfolding event and its impact on memory relied on the disruption of neural stability patterns preceded by the neural reactivation of the memory representations of the just-encoded event. Our study includes data from 3 different experiments whereby participants encoded images of target objects preceded by trial-unique sequences of events depicting daily routine. We found that neural stability patterns gradually increased throughout the ongoing exposure to a schema-consistent episodic narrative and that the brain stability pattern was interrupted when the encoding of an object of the event was lowly congruent within the ongoing schema representation. We found that the decrease in neural stability for low-congruence items was best seen at ∼1000 ms from object encoding onset when compared to high-congruence items and that this effect was preceded by an enhanced N400 ERP and an increased degree of neural reactivation of the just-encoded episode for low-congruence items. Current results offer new insights into the neural mechanisms and their temporal orchestration that are engaged during online encoding of schema-consistent episodic narratives and the detection of incongruencies.

## Introduction

Experience is guided by internal representation models of the environment, or knowledge schemas, with an impact on perception and memory (Gilboa and Marlatte, 2017). Schemas are thought to be central in organising our everyday life behaviour by giving stability and predictability to the structure of the world (Gershman et al., 2014). Thus, despite the ever-changing sequence of inputs of our experience, schemas bring relatedness and comprehension of unfolding events by anticipating stereotyped or congruent-like elements to encounter next. A computational advantage to the memory systems is, therefore, that schema-consistent items can be added to an existing schema without requiring alterations or extensions to it (McClelland et al., 2020). Accordingly, when elements of the unfolding experience are congruent with expected representations from a currently activated schema, they are rapidly integrated into the memory model of the event (van Kesteren et al., 2007; McClelland et al., 2020; Tse et al., 2007 and 2011). However, when our predictions are incorrect, we must update our internal models of the world to support adaptive behaviour. Nevertheless, the neural mechanisms that support memory integration and updating of an unfolding event remain unclear.

If an internal memory representation is stable over time, then some properties of its underlying neural implementations may also exhibit invariance during encoding. Indeed, neuroimaging studies in humans observed stable brain patterns of activity during the encoding of continuous streams of audio-visual inputs and that shifts in neural stability are coincident with the detection of unexpected elements in the unfolding stream (i.e., event boundaries) (Baldassano et al., 2017). Similarly, Sinclair et al. (2021) recently showed that hippocampal activation patterns stabilised during the encoding of a narrative episode, akin to sustained representations accumulated during an unfolding schema-congruent event. Intriguingly, this study also revealed that when the narrative was suddenly interrupted, the ongoing stability of the neural activity became disrupted, reflecting the need to update the sustained representation of the event model.

However, the notion that the detection of incongruencies of an unfolding event engenders a disruption of the ongoing representation challenges a set of findings that found no memory disturbance or even improvement for surprising events (e.g., Greve et al., 2017; Greve et al., 2019; Quent et al., 2021; Frank et al., 2018, 2020; Rouhani et al., 2018; Chen et al., 2015; Pine et al., 2018). This literature relies on the idea that mnemonic prediction error enhances hippocampal biases toward encoding (van Kesteren et al., 2017; Bein et al., 2020) and that this shift in encoding strategy reflects the need to evaluate and, if necessary, update the representational content of the ongoing experience with the current incongruent event.

How does the brain accommodate these two seemingly opposite lines of research evidence, namely, that mnemonic prediction errors disrupt ongoing neural representations of unfolding events and, at the same time, promote the update of the ongoing memory model during encoding? Here, we asked whether this process is supported by distinct brain mechanisms that occur rapidly (in the order of milliseconds) but are sequentially orchestrated over time. More specifically, drawing on past theoretical (McClelland et al., 2020) and empirical research (Sols et al., 2017; Silva et al., 2019; Wu et al., 2021), we hypothesise that the subjective degree of an item’s congruence with an unfolding experience is determined by an evaluation process guided by a rapid reactivation of the encoded event. The concomitant representation of the new element and the reactivated memory of the just-encoded event would promote the effective and rapid assessment of the extent to which the novel element matches or mismatches expectations driven by the unfolding event. As a result, the brain would be able to either assimilate the new item with the ongoing memory representation by preserving a stable state of neural pattern of activity or, alternatively, disrupt it to promote its update.

To test this hypothesis, we recorded scalp electrophysiological (EEG) activity while healthy participants encoded images of target objects preceded by trial-unique sequences of four pictures of events depicting a routine from everyday life (**Figure 1a**). The sequence of pictures preceding the target object image was thought to mimic a realistic unfolding episodic event with the aim to provide a gradual schema-consistent narrative that determined whether specific target objects matched or mismatched expected occurrences within that context. Importantly, participants were instructed to rate the perceived congruence of the item in relation to the previously encoded event sequence episode, thereby allowing us to assess the degree of perceived congruence of the target object for every single trial at an individual level. To examine how object congruence shaped memory for the target object, a surprise recognition memory test was administered to the participants after the encoding phase. In experiment 1, we first asked participants to indicate whether a label word referred to object pictures encoded in the previous phase, and if so, to recognize which of two very similar pictures was the exact one presented during the encoding phase (**Figure 1b**). This later test allowed assessment of the extent to which encoding congruence detailed memory representation (e.g., Bein et al., 2020). We ran two additional follow-up behavioural experiments on a separate sample of participants to further scrutinise congruence-shaped long-term memory. The two experiments consisted of a similar structure and materials used in Experiment 1 but differed in the format of the recognition phase. In Experiment 2, we presented pictures that corresponded to the ones shown during the encoding phase and pictures that depicted similar objects but differed in small visual details (lures), thereby allowing us to assess whether possible differences in memory as a function of encoded congruence were independent of the format of the test (**Figure 1c**). In experiment 3, we asked participants to indicate whether object pictures were encoded together with a selected picture of the episodic sequence (**Figure 1d**). This test allowed examining whether the perceived congruence of the object within an episodic narrative influenced how the two become associated in long-term memory.

**Figure 1.**
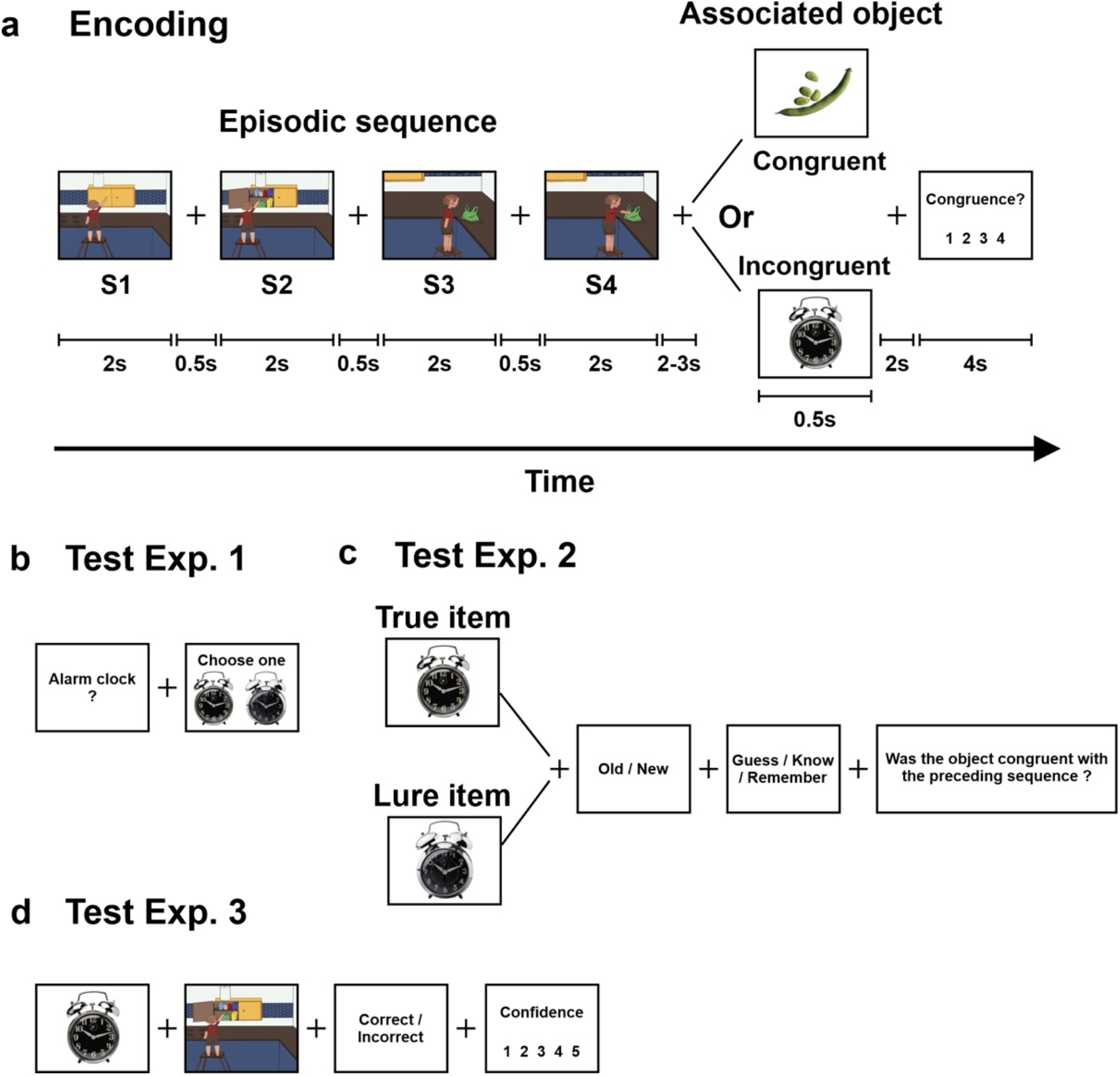
Encoding and recognition memory task design for Experiments 1, 2 and 3. **(a)** During the encoding phase for all experiments, participants encoded episodic sequences composed of 4 photographs depicting a routine domestic episode. These were followed by highly congruent or low congruent/incongruent object pictures. Participants indicated the degree of congruence between the episode and the object. **(b)** In Experiment 1, memory of object pictures was tested by the object word label followed by a true and a lure item of the same object. Participants had to indicate the correct picture presented during the encoding phase. **(c)** In Experiment 2, memory of the encoded objects was assessed by requesting participants to discriminate whether true or lure items corresponded to those presented during the encoding phase. This was followed by a ‘guess/know/remember’ task and ended with asking the participants to indicate whether the picture object was encoded within a congruent episodic event. **(d)** In Experiment 3, each object picture presented during the encoding phase was displayed together with one image from an episodic sequence. Participants were required to indicate whether the object and episodic image picture corresponded with the episodic + object picture presented in the same trial during the encoding phase. (a) and (d) depict drawings of episodes instead of pictures of real-life events due to bioRxiv policy on not displaying pictures of real people.

## Material and Methods

### Participants

Participants were healthy college students from the University of Barcelona who had normal or corrected-to-normal vision and reported no history of medical, neurological or psychiatric disorders. Thirty-three participants (26 females, M = 20.94 years, SD = 3.24 years) were recruited and were paid €10/h for their participation in Experiment 1. Four participants were excluded due to loss of EEG data for technical reasons. Thirteen (11 female, M = 22.17 years, SD = 2.33) and eighteen (16 female, M = 23.05 years, SD = 6.55) participants were recruited and paid €5/hour for their participation in the follow up Experiment 2 and 3, respectively. All participants signed informed consent, approved by the University of Barcelona Ethics Committee.

### Stimuli

Experimental stimuli consisted of 160 photographs of household objects and 80 episodic sequences, each comprising 4 photographs. There were 80 different household objects included, each with two slightly different versions, for a total of 160 photographs. Episodic sequences consisted of 4 snapshots in temporal order depicting a person moving around and interacting with the surroundings in different house rooms during a short period of time. Each sequential episode was designed to match one of the 80 household objects. The object images were taken from the Stark lab set of stimuli, freely available at (http://faculty.sites.uci.edu/starklab/mnemonic-similarity-task-mst/). The pictures of the episodic sequences did not actually contain the matching household object. Instead, the sequences were designed in a way that the matching object could fit in or make sense within the given sequence. In other words, the matching object could be expected to be encountered in the situation depicted in the episodic sequence. Each episodic sequence was designed to be congruent with its specific corresponding (congruent) object.

### Experimental design

The experiment design of the three studies consisted of an encoding phase and a test phase. The encoding phase was the same in all three studies. Participants were presented in trial with an episodic sequence followed by a picture object. Participants were asked to rate the degree of congruence on a scale from 1 (i.e., does not fit in) to 4 (i.e., fits in very well) of the target picture object in relation to the context formed by the succession of the 4 preceding episodic sequence images (**Figure 1a**). The encoding phase included a total of 80 trials, each consisting of an episodic sequence followed by an object. Two versions of the encoding phase were constructed so that 40 of the episodic sequences could be perceived as highly congruent with picture objects by the participant. This yielded a total of an *a priori* possible 40 high and 40 low congruent sequence–object pairings. The order of the trials at encoding was randomised for each participant.

Each trial started with the appearance of a fixation cross on the screen for a random duration of 2000 to 4000 ms. Afterwards, an episodic sequence consisting of four photographs was presented. Each of the four photographs was presented on a white background for 2000 ms, one at a time in temporal order, separated by the presentation of a fixation cross for 500 ms. After the episodic sequence was presented, a fixation cross appeared on the screen for 2000 to 3000 ms, separating the episodic sequence from the presentation of the following object. The picture of the object was presented on a white background for 500 ms, followed by the appearance of a fixation cross for 2 seconds. Finally, a screen was presented with the word ‘Congruence?’ and the digits ‘1-2-3-4’ below, upon which participants had to indicate, within a maximum of 4 seconds, the degree of congruence between the object and the just-encoded episodic sequence by pressing 1, 2, 3 or 4 in the keyboard. Participants were previously instructed to respond thoughtfully as quickly and accurately as possible. As soon as they responded, a fixation cross was presented for 500 ms, and the next trial began. The encoding phase lasted around 30 min. After the 80 episodic sequences and objects were presented, the encoding phase was finished.

The encoding phase was followed by a ∼10 min interference task consisting of choosing the correct answer to simple additions and subtractions that appeared on the screen. Participants were told to respond as quickly as possible, although no time limit was imposed. The distraction task ensured the participants would not rehearse the pictures they had previously seen.

### Recognition memory test

After a break of ∼10 minutes, a recognition memory test was presented unexpectedly to the participants in the three studies.

In Experiment 1, the test included 160 object words and 160 object picture pairs of each word. Eighty words and object pictures corresponded to previously presented objects in the encoding phase (‘Old’), whereas the other 80 were non-related (‘New’) objects (**Figure 1b**). Each picture pair depicted the same object but with small changes in specific features between each other (e.g., orientation, colour, etc.). Each trial began with a fixation cross lasting from 3 to 5 seconds at random. Subsequently, each word was presented for a maximum of 6 seconds with a question mark below ‘?’. Participants were instructed to press ‘1’ on the keyboard if they thought the word referred to an object presented during encoding (Old) or ‘2’ if not (New). If the participant responded ‘Old’ to a word, then two pictures of the object word were presented on the screen for a maximum of 6 seconds (**Figure 1b**). Pictures were presented on the computer screen, with one item to the left and one to the right of fixation. The left/right assignment was randomly chosen in each trial. The two pictures from each pair were almost identical, but only one corresponded to the exact one presented in the encoding phase (true), whereas the other one served as a lure. For example, if the participant had seen the photograph of a hammer and later, during the test, correctly identified the word ‘hammer’ as one of the objects she saw, then two photographs of similar-looking hammers appeared. The participant was instructed to identify the picture object that was exactly like the one presented in the encoding phase by pressing 1 if it was the left photograph or 2 if it was the right one. The picture pair presented when a participant misclassified as ‘Old’ a new word was almost identical, though neither of the two pictures had been seen in the encoding phase. The order of the presentation of word + picture pairs in the test was randomised before each participant started the test. The recognition memory test lasted ∼20 min.

In Experiment 2, memory of the encoded objects was assessed by asking participants to recognise them in a set of pictures presented randomly in the test phase. The test included 40 target objects previously presented at encoding (True items) and 40 objects highly similar to the target objects presented at encoding but with some changes in their specific features (e.g., orientation, colour, etc.) (Lure items) (**Figure 1c**). In total, the recognition memory test comprised 80 items. 40 of the picture objects (20 True and 20 Lure) related to pictures encoded in high-congruent trials, and the other 40 picture objects (20 True and 20 Lure) related to pictures encoded in low-congruent trials at the encoding phase. Two versions of the test phase counterbalancing True/Lure and high/low congruency conditions were prepared and assigned randomly to the participant’s sample. Each trial began with a fixation cross lasting from 3 to 5 seconds at random. One object picture then remained on the screen for a maximum of 8 seconds with the question ‘Did this object appear before?’, and participants had to indicate on a keyboard whether the same item was presented during encoding (‘1’ – Old and ‘2’ – New). Participants were told only items the same as items presented during encoding were correct answers. ‘Old’ responses were followed first by a ‘guess/know/remember’ judgment of the picture and later by a question referring to the semantic context: ‘Was this object encountered in a high/low congruent context?’. Participants had a maximum of 8 seconds to respond to each question.

In the test phase of Experiment 3, the 80 object pictures presented during the encoding phase were included in the test. In each trial, each object picture was presented together with the first or the second image from each of the episodic sequences presented in the encoding phase (**Figure 1d**). Participants were requested to indicate whether the object picture and episodic image picture matched the episodic sequence and object picture presented during the encoding phase. Half of the object+episode picture pairs presented in the test matched the encoding ones, whereas the others were randomly paired with each other. The total set of 80 picture pairs was constructed so that 40 old ones included 20 object+episode pairs encoded in the high-congruency condition and 20 in the low-congruency condition. The same distribution pattern was used to construct the set of object+episode new picture pairings (i.e., those that do not match the trials presented at encoding). Two different versions of 40 old and 40 new sets of picture pairings were created by controlling so that in one version, a picture object was paired with a matched image from the encoded episodic sequence (Old) and to an unmatched image (New) in the other. The two versions were assigned randomly to the participant’s sample. Each trial began with a fixation cross lasting from 3 to 5 seconds at random. The object and the episodic sequence picture were then presented on the screen. Participants were instructed to answer whether both photographs had been presented together in the same trial during the encoding phase (by pressing ‘1’ on the keyboard) or not (by pressing ‘2’). Participants were asked to rate their confidence in their previous response from 1 (‘no confidence’) to 5 (‘absolute confidence’) with the same numbers on the keyboard. Object and episodic pictures were presented on the computer screen, with one item to the left and one to the right of fixation. The left/right assignment was randomly chosen on each trial.

### Behavioural data analyses

Paired Student t-test was used to compare participants’ performance (measured in percentage) between conditions. Repeated measures ANOVA was used to statistically assess differences between participants’ performance when they included more than two variables. Statistical significance threshold was set at *p* < 0.05.

### EEG recording and preprocessing

In study 1, EEG was recorded during the task using a 32-channel system at a sampling rate of 500 Hz, with an online band-pass filter set at 0.01-100 Hz, using a BrainAmp amplifier and tin electrodes mounted in an electrocap (Electro-Cap International) located at 29 standard positions (Fp1/2, Fz, F7/8, F3/4, Fcz, Fc1/2, Fc5/6, Cz, C3/4, T3/4, Cp1/2, Cp5/6, Pz, P3/4, T5/6, Po1/2, Oz) and at the left and right mastoids. An electrode placed at the lateral outer canthus of the right eye served as an online reference. EEG was re-referenced offline to the average of all channels. Vertical eye movements were monitored with an electrode at the infraorbital ridge of the right eye. Electrode impedances were kept below 5 kΩ. EEG was low-pass filtered offline at 30 Hz. We applied the Parks-McClellan Notch filter using the toolbox ERPLAB (http://erpinfo.org/erplab).

For each participant, we extracted the EEG epochs for each encoding image. Epochs had a duration of 2000 ms for images from the episodic sequence and 2500ms for images of the picture object and were baseline corrected to the pre-stimulus interval (−100 to 0 ms). Epochs with maximum absolute amplitude over 100 µV were discarded for further analysis. For later analysis, all the epochs were smoothed by averaging data via a moving window of 100 ms (excluding the baseline period) and then downsampled by a factor of 5.

### Neural stability analysis

To account for whether the ongoing encoding of an episodic sequence of pictures elicited a gradual increase in stable brain activity patterns, we implemented a temporally resolved similarity analysis using Pearson correlation coefficients, which are insensitive to the absolute amplitude and variance of the EEG response. The correlation analysis on EEG data was made at the individual level and to each time point separately and included spatial (i.e., scalp voltages from all the 29 electrodes) features of the resulting EEG single trials. To examine how a schema-consistent sequence’s unfolding modulated the stabilisation of activity patterns, we correlated the EEG patterns of activity elicited by the 1^st^ and the 2^nd^ pictures and compared them to the correlation obtained by the 3^rd^ and the 4^th^ picture of the episodic sequence. This analysis was computed at the trial level by randomly creating pairs of correlation analyses from pictures from different episodic sequences. For each participant, we created 200 permutations of possible unique pairings, and the final cross-correlation output resulted from averaging the point-to-point Fisher’s z scores correlation values across the 200 permutations.

A similar analysis was performed on EEG patterns elicited by picture objects. For each participant, we randomly created 200 sets of unique picture pairs that the participant rated as High congruent (i.e., rated as 1 or 2 during the encoding phase) or Low congruent (i.e., rated as 3 or 4 during the encoding phase).

We implemented a cluster-based permutation test to account for neural stability differences between picture order within the episodic sequence and between High and Low congruence conditions (Maris and Oostenveld, 2007). It identifies clusters of significant points in the resulting 1D matrix in a data-driven manner and addresses the multiple-comparison problem by using a nonparametric statistical method based on cluster-level randomisation testing to control the family-wise error rate. Statistics were computed for every time point, and the time points whose statistical values were larger than a threshold (*p* < 0.05, two-tail) were selected and clustered into connected sets based on adjacency points in the 1D matrix. The observed cluster-level statistics were calculated by taking the sum of the statistical values within a cluster. Condition labels were then permuted 1000 times to simulate the null hypothesis, and the maximum cluster statistic was chosen to construct a distribution of the cluster-level statistics under the null hypothesis. The nonparametric statistical test was obtained by calculating the proportion of randomised test statistics that exceeded the observed cluster-level statistics.

### Representational Similarity Analysis (RSA)

RSA was performed timepoint-to-timepoint at trial level and upon spatial features (i.e., scalp voltages from all the 29 electrodes) (Silva et al., 2017; Wu et al., 2021). RSA was conducted between the EEG signal of each encoding image (i.e., the image at 1^st^, 2^nd^, 3^rd^, 4^th^ position in a sequence) with the EEG signal of the corresponding offset period (i.e., the matching object with the sequence and the followed fixation cross). Point-to-point Pearson correlation values were then calculated, resulting in a 2D similarity matrix with the size of 200×250 (each time point represents 10 ms, given the 500 Hz EEG recording sampling rate and a down-sampling factor of 5). The *x*-axis of the matrix represented the object and offset time points, and the *y*-axis represented the time points of sequence picture encoding. The output 2D matrix represents the overall degree of neural pattern similarity between EEG elicited for each pair of encoding images and its corresponding sequence offset.

### Linear-mixed effect model

To further explore in the trial base how the neural pattern was associated with the subjective feeling of the congruence and its impact on subsequent memory of the object image, we applied a Linear Mixed-Effect Model (LMM) on ERPs elicited during the offset period as well as the pattern similarity between encoding and offset.

For offset ERPs, we first grouped 29 electrodes into 6 scalp regions. To obtain more stable spatial patterns, border electrodes between regions were included in each neighbouring region (Lu et al., 2015; Sols et al., 2017). As a result, the 6 regions were defined as follows: region 1 (FP1, Fz, F3, F7, FCz, FC1, FC5); region 2 (FP2, Fz, F4, FCz, FC2, FC6); region 3 (FCz, FC1, FC5, Cz, C3, T3, Cp1, Cp5); region 4 (FCz, FC2, FC6, Cz, C4, T4, CP2, CP6); region 5 (CP1, CP5, Pz, P3, T5, PO1, Oz); region 6 (CP2, CP6, Pz, P4, T6, PO2, Oz). The LMM was constructed separately for each region. ERPs voltages at each offset time point were averaged across electrodes within the region and then introduced in our LMM as the dependent variable. The rating of the object’s congruence to the sequences (on a scale of 1 to 4) and the memory for the object (whether the word and the image were correctly recognised during the test) were included in the model as fixed effect variables. The subject was then introduced into the model as the grouping variable, with random intercept and a fixed slope for each fixed effect variable. The statistical significance was adjusted by Bonferroni correction for each fixed effect variable taking into account both the number of regions and the number of time points, resulting in a corrected alpha level of α = 3.33×10^−5^ (0.05/(6*250)).

We also applied LMM on output from RSA. We identified on the resulting 2D similarity matrix the time point of encoding and offset where the pattern similarity reached the peak value. We then averaged at a single-trial level the similarity value across ± 50 ms (11 data points) around the peak encoding time point for each offset time point, resulting in a 1D similarity value with the length of 250 time points, covering the whole offset period. For each time point, we constructed the LMM with similarity value as the dependent variable. Then 3 fixed-effect variables were introduced into the model: the image position in the sequence (1^st^, 2^nd^, 3^rd^, and 4^th^), congruence rating and memory for the object. Subject was included in the model as the grouping variable, with random intercept and a fixed slope for each fixed-effect variable. The statistical significance for each fixed-effect variable was Bonferroni corrected with a thresholded alpha level of α = 2×10^−4^ (0.05/250).

## Results

### Behavioural results from Experiment 1

In Experiment 1, the proportion of trials rated as highly congruent (‘1’ or ‘2’) was 45.17% (SD = 8.59%) and 15.43% (SD = 6.42%), respectively, and lowly congruent (‘3’ or ‘4’), respectively, was 13.45% (SD = 6.36%) and 25.39% (SD = 7.47%) with the episode.

In general, participants were accurate in recognizing words referring to picture objects learnt during the encoding phase (Mean = 71.09%, SD = 12.80%; above chance: *t*(28) = 8.41, *p* < 0.001). They showed greater accuracy for words related to picture objects encoded with high (Mean = 76.07%, SD = 13.27%) than low (Mean = 66.11%, SD = 13.89%) congruency to the episodic context during the encoding phase (paired t-test: *t*_(28)_ = 5.89, *p* < 0.001) (**Figure 2a**). For those correctly recognised words, in the next picture recognition test, participants were accurate in correctly identifying the exact object picture presented in the encoding phase (Mean = 73.13%, SD = 8.02%; above chance: *t*_(28)_ = 15.53, *p* < 0.001). They performed similarly for pictures rated as high (Mean = 74.61%, SD = 11.63%) and low (Mean = 72.61%, SD = 8.61%) congruency to the episodic context (paired *t*-test: *t*_(28)_ = 0.39, *p* = 0.878) (**Figure 2b**).

**Figure 2.**
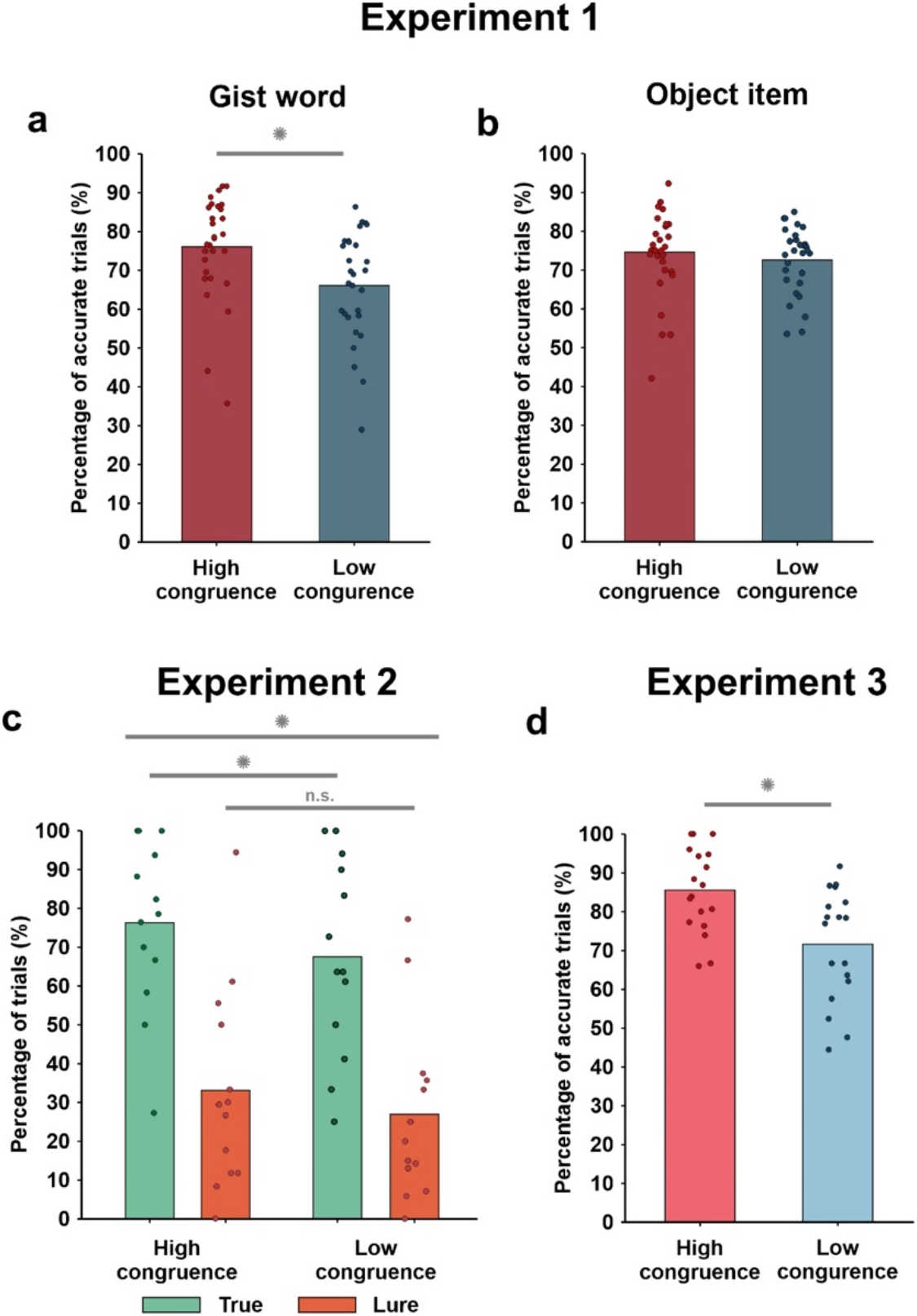
Behavioural results of Experiments 1, 2 and 3. **(a)** Participants’ test accuracy on recognising the gist word separated by whether the picture object was perceived by the participant as highly or lowly congruent with the episodic sequence during encoding in Experiment 1. **(b)** Participants’ test accuracy on identifying the exact object picture presented in the encoding phase after correctly recognising the ‘Old’ gist word. The accuracy was separated by whether the picture object was perceived by the participant as highly or lowly congruent with the episodic sequence during encoding in Experiment 1. **(c)** Participants’ memory accuracy (in percentage) at the recognition test of Experiment 2 for True (hit) and Lure (correct rejection) as a function of whether the picture object was perceived by the participant as having high or low congruence with the episodic sequence during encoding. **(d)** Participants’ memory accuracy (in percentage) in the recognition memory test of Experiment 3. In all plots, dots represent values for an individual subject. * indicates *p* < 0.05. ‘n.s.’ indicates *p* >0.05.

To specify the subsequent memory strength for objects, for all later analyses, we classified the memory performance of a trial based on whether the object image was correctly recognised during the test. Therefore, successful recognition of both gist word and object image was considered as a ‘remembered item’ condition, while either failing in recognition of gist word or image was considered as a ‘forgotten item’ condition. This separation also rendered a balanced percentage of trials across conditions, with 53.53% (SD = 11.43%) for ‘remembered’ condition and 46.47% (SD = 11.43%) for ‘forgotten’ condition (Wilcoxon signed-rank test: *z* = 1.74, *p* = 0.08).

### Behavioural results from Experiments 2 and 3

The results of Experiment 1 revealed that memories of objects that were perceived as highly congruent with the episodic context were later better remembered. However, the results of this study relayed on a test that changed the perceptual format at test, that is, objects were encoded as a picture and tested as a word at test, which lacks the visual details of the picture object and engenders semantic processing. A concern of this change could be that memory congruency benefits were to some extent explained by differential processes taking place during retrieval. Information that is highly congruent with prior knowledge is often found to be better remembered than low congruent information, putatively because of an increase in semantic associations and relational integration (Staresina et al., 2009; Atienza et al., 2011; van Kesteren et al., 2014; Bein et al., 2015; also see Craik and Tulving, 1975). Experiment 2 addressed this concern by testing object memories in the same visual format depicted at the encoding phase.

In Experiment 2, the proportion of trials rated as high (‘4’) and low congruent (‘1’) were 50.10% (SD = 10.58%) and 42.21% (SD = 4.98%), respectively, and the average proportion of intermediate levels of congruency was very low (‘2’: Mean = 2.69%, SD = 3.42%; and ‘3’: Mean = 5%, SD = 5.68%). A comparison of the proportion of trials rated as high congruent (‘4’) and low congruent (‘1’) revealed they were not significantly different (t_(12)_ = 2.03, *p* = 0.06). Given the low proportion of trials rated by the participants with intermediate level of congruence (i.e., < 10% on average), and to ensure a proper orthogonalization of possible memory effects at the test driven by encoding congruency, we included in the subsequent analyses trials rated as high (‘4’) or low congruent (‘1’) by the participants.

In general, participants were highly accurate in recognizing True Old object pictures that were encoded on a highly congruent (Mean = 0.99, SD = 0.01) and on a low-congruence episodic context (Mean = 1, SD = 0). They also showed to be prone to misclassify Lure items as Old in the two encoding conditions (High congruent: Mean: 0.48, SD = 0.23; Low congruent: Mean = 0.47, SD = 0.22). To investigate whether retrieval accuracy differed as a function of encoding congruence, we applied a repeated measures ANOVA, including picture type (true vs. lure), congruence (High vs Low congruent) and subjective retrieval quality (remember vs. know/guess), as within-subject factors. The results confirmed that participants were more accurate in correctly recognizing True items than rejecting Lure items as Old (main effect of picture type: *F*_(1,12)_ = 81.76, *p* < 0.001) but, as expected, participants differed in their accuracy for True and Lure items as a function of whether the retrieval was catalogued as ‘remember’ or ‘guess/know’ (main effect of remember: *F*_(1,12)_ = 7.88, *p* = 0.016 and an interaction picture type x remember: (*F*_(1,12)_ = 16.97, *p* = 0.001). However, we did not find a main effect of congruence (*F*_(1,12)_ = 0.35, *p* = 0.85) or a congruence x item type interaction (*F*_(1,12)_ = 0.16, *p* = 0.67), but found a significant retrieval quality x congruence interaction (*F*_(1,12)_ = 6.95, *p* = 0.02), which suggested that congruence at encoding may affect retrieval accuracy only for when participants were capable to retrieve object information as a function of the ability to retrieve their context (**Figure 2c**). To unpack this finding, we ran separate repeated measures ANOVA, including item type and congruence as within-subject factors, for participants’ accuracy as a function subjective retrieval quality. The results indicated that participants were more accurate in correctly identifying True items than in misclassifying as Old the Lure items in the remember condition (main effect of item type: *F*_(1,12)_ = 103.22, *p* < 0.001) and that participants’ accuracy differed for high and low congruent encoded items (main effect of congruence: *F*_(1,12)_ = 9.52, *p* = 0.009; congruence x item type interaction: *F*_(1,12)_ = 0.44, *p* = 0.52). A post-hoc paired student t-test indicated higher accuracy for high than low congruent True items (*t*_(12)_ = 2.65, *p* = 0.02) and a trend towards statistical significance for Lure items (*t*_(12)_ = 2.07, *p* = 0.06). The ANOVA on response accuracy for ‘guess/know’ condition showed a similar trend directions in the effects, though none of the effects reached statistical significance (main effect of item type: *F*_(1,12)_ = 4.17, *p* = 0.06; main effect of congruency: *F*_(1,12)_ = 3.59, *p* = 0.08; item type x congruency interaction: *F*_(1,12)_ = 0.78, *p* = 0.39). Collectively, the results of Experiment 2 replicated the effects observed in Experiment 1, by showing that retrieval accuracy was higher for items that were preceded by a high than a low congruent episodic context with the object. However, they also reveal that items encoded in a high congruent context are more prone to errors at retrieval, suggesting the possibility that the benefit of congruence may also come with a decreased detailed item representation in long-term memory.

We next sought to examine whether the retrieval benefits of encoding congruence observed in Experiments 1 and 2 were also accompanied by an increased degree of binding of the object to the preceding episodic context. Thus, in Experiment 3, participants were required to judge whether pairs of pictures, including one of the encoded objects and one picture from the sequence episodes, were seen together or not during encoding. As in study 1 and 2, the proportion of trials rated as high (‘4’) and low congruent (‘1’) were 35.39% (SD = 12.63%) and 37.67% (SD = 10.06%), respectively, and the average proportion of intermediate levels of congruency was much lower (‘2’: Mean = 12.92%, SD = 9.97%; and ‘3’: Mean = 10.81%, SD = 5.68%). A comparison of the proportion of trials rated as high congruent (‘1’) and low congruent (‘4’) revealed they were not statistically different (*t*_(17)_ = −0.85, *p* = 0.41). Therefore, as in study 2, we analysed memory accuracy for items classified as ‘1’ or ‘4’ at encoding. The results of this study revealed that participants were accurate in correctly identifying the encoded correspondence between objects and the episodic context (Mean = 0.79, SD = 0.12), but that the accuracy was higher (paired student *t*-test, *t*_(17)_ = 5.59, *p* < 0.001) and more confident (*t*_(17)_ = 6.78, *p* < 0.001) when the association between the object and the context at encoding was high rather than low congruent (**Figure 2d**).

### Neural stability and episodic congruence

We first examined whether EEG states of stability gradually increased throughout the encoding of a schema consistent images of the narrative sequence. We hypothesised that schema consistency for picture sequences would promote neural stability, and that this would be observed as a gradual increase in across-item EEG pattern similarity (**Figure 3a**) of the EEG patterns elicited throughout the picture sequence. To address this issue, we calculated a time-resolved neural similarity analysis between EEG patterns elicited by initial (i.e., 1^st^ and 2^nd^) and final (3^rd^ and 4^th^) picture items across the sequences. Confirming our hypothesis, we found that the 3^rd^ and the 4^th^ picture elicited more stable patterns of EEG activity from 1030 ms to 2500 ms from picture onset (Cluster statistics: *p* = 0.009, corrected for multiple comparisons, sum *T*-values = 259.78) (**Figure 3b**).

**Figure 3.**
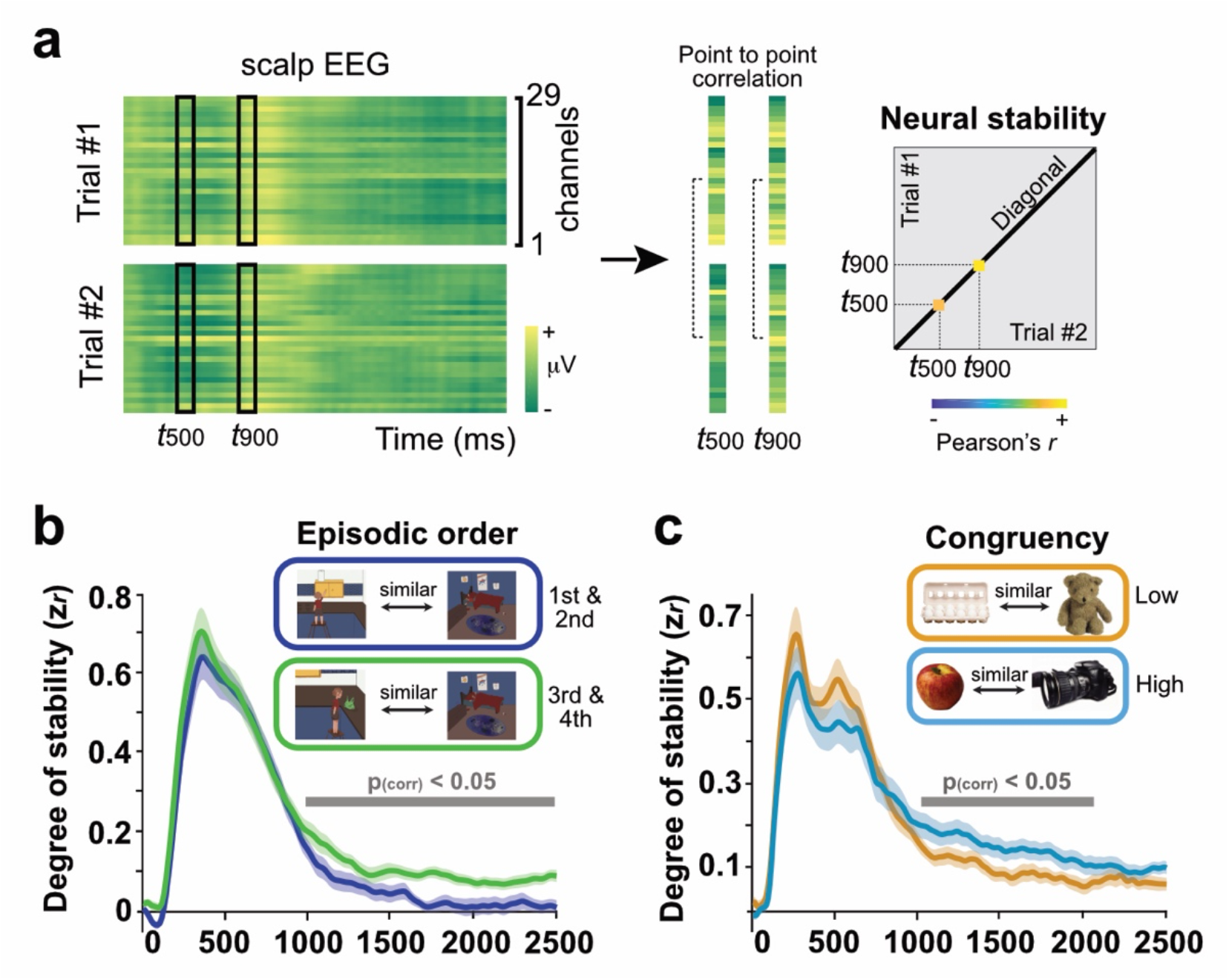
Neural stability and congruence. **(a)** Schematic representation of the analysis. A temporal cross-stimuli correlation matrix is generated from the EEG data for each participant. **(b)** Point-to-point participants’ average degree of neural stability between 1^st^ and 2^nd^ and 3^rd^ and 4^th^ picture across different episodic sequences. **(c)** Point-to-point participants’ average degree of neural stability across object pictures rated lowly and highly congruent with the preceding episodic event at encoding. The shaded area represents standard error (SEM) across subjects. The thick grey line depicts the timing of the significant cluster between conditions (*p* < 0.05, cluster-based permutation test). (d) depicts drawings of episodic instances instead of pictures of real-life events due to bioRxiv policy on not displaying pictures of real people.

Next, we evaluated whether EEG states of stability were associated with the encoding of object pictures that were highly or lowly congruent with the preceding episode. We hypothesised that congruency induced higher states of stability and that this elevated state of neural stability would be reflected as an increase in neural similarity upon object encoding, rendering them more similar than objects perceived as lowly congruent with the preceding episodic context. To address this issue, we compared the cross-temporal correlation analysis to EEG activity patterns from different pictures within each condition. We reasoned that if congruency induced patterns of neural stability, this should be reflected as a persistent general pattern of activity during encoding, independently of the depicted object. Please note that the assignment of episode-object associations was counterbalanced between participants and, therefore, an object encoded highly congruently with an episode in one participant was encoded lowly congruently by another participant. The results of this analysis showed that high-congruence items elicited more stable patterns of EEG activity than low-congruence items from 1036 ms to 2008 ms from picture onset (Cluster statistics: *p* = 0.02, corrected for multiple comparisons, sum *T*-values = 452.90) (**Figure 3c**). These results indicate that context-dependent neural state of stability is modulated by episodic congruency at encoding.

### The N400 signals low congruent items from encoding episodes

We hypothesised that the disruption of neural stability elicited by the detection of low-congruence objects from the unfolding schema-consistent episodic narrative would be preceded by a prediction error signal in the brain. We aimed to identify such an error signal in the EEG by means of a transient increase in the N400 ERP component, which has been widely related to incongruence detection in the literature (Kutas and Federmeier, 2008). To assess this issue in our data, we grouped the 29 electrodes into 6 regions and averaged the epochs across electrodes within each region. We then introduced the voltage value into the LMM as the dependent variable and included participants’ ability to correctly recognise the picture image at test congruence rating provided at encoding as the main fixed-effect variables. Subject was included as the grouping variable. This analysis was conducted for each time point and each scalp region separately.

The results of the LMM analysis revealed significant effects at specific scalp regions and temporal points for both memory and congruence. For memory, later forgotten objects elicited significantly more negative ERPs amplitude at 520 ms to 870 ms from picture onset distributed over the frontal scalp region. For congruence, we found more substantial negative amplitude with objects rated as less congruent with the preceding episodic context. This effect was significant from 410 ms to 730 ms from picture onset and was distributed over frontocentral scalp regions (**Figure 4a**).

**Figure 4.**
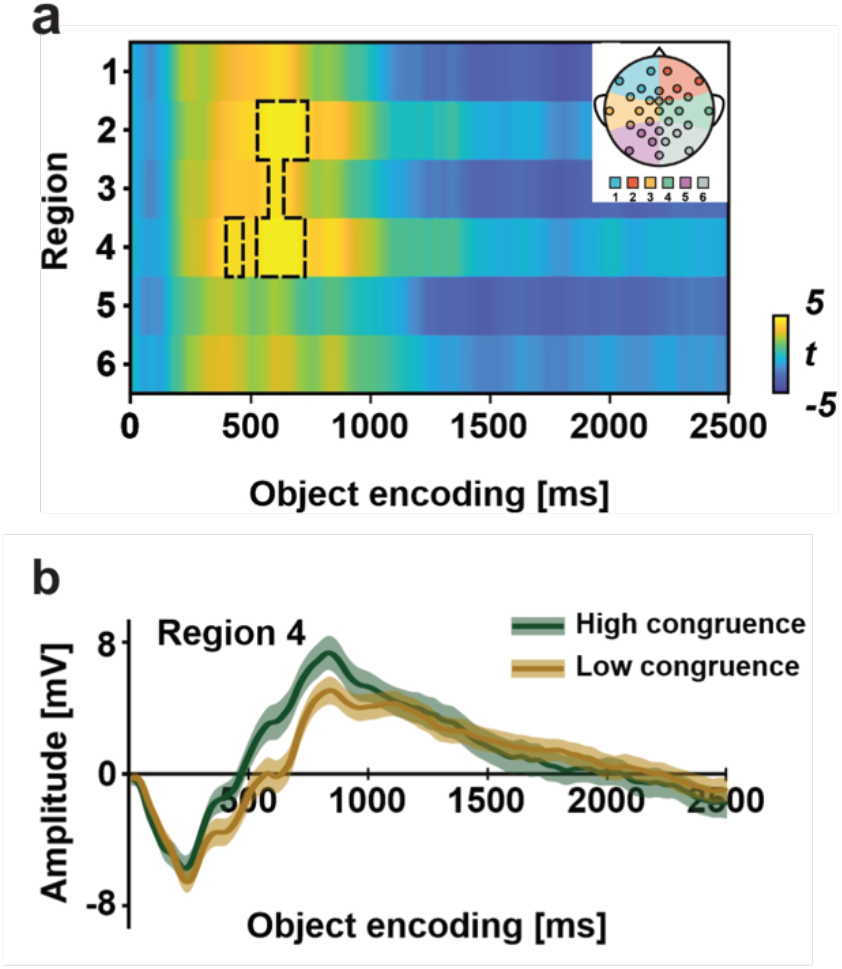
LMM on ERPs elicited by object pictures. 29 electrodes were grouped into 6 scalp regions, and boundary electrodes were included in either neighbouring region to obtain more stable spatial patterns. **(a)** *t*-value resulting from the LMM analysis at each region and time point as a function of congruence. **(b)** Participants’ averaged ERPs from representative region 4 (note that for visual illustration, trials with congruence ratings of 1 and 2 were grouped and averaged as Low congruence and trials with congruence ratings of 3 and 4 were grouped as High congruence). The shaded area represents standard error (SEM) across subjects. A black dashed line on the statistical map marks the area where the *t*-statistics exceed the significance threshold (*p* < 0.05) with the alpha level adjusted with Bonferroni correction.

### Neural reinstatement induced by low contextual congruency

We next sought to test that prediction error elicited by low-congruence items would be accompanied by more robust reactivation of the just-encoded episodic elements. To address this issue, we implemented a trial-based and temporally resolved neural similarity analysis between EEG data elicited at picture object and EEG data elicited by each image of the preceding episodic context. The results of this approach revealed an increase in neural similarity between EEG patterns from ∼100ms - 700ms at object picture onset and EEG patterns of activity between ∼100ms - 750ms from the onset of pictures within the episodic sequence (**Figure 5a**).

**Figure 5.**
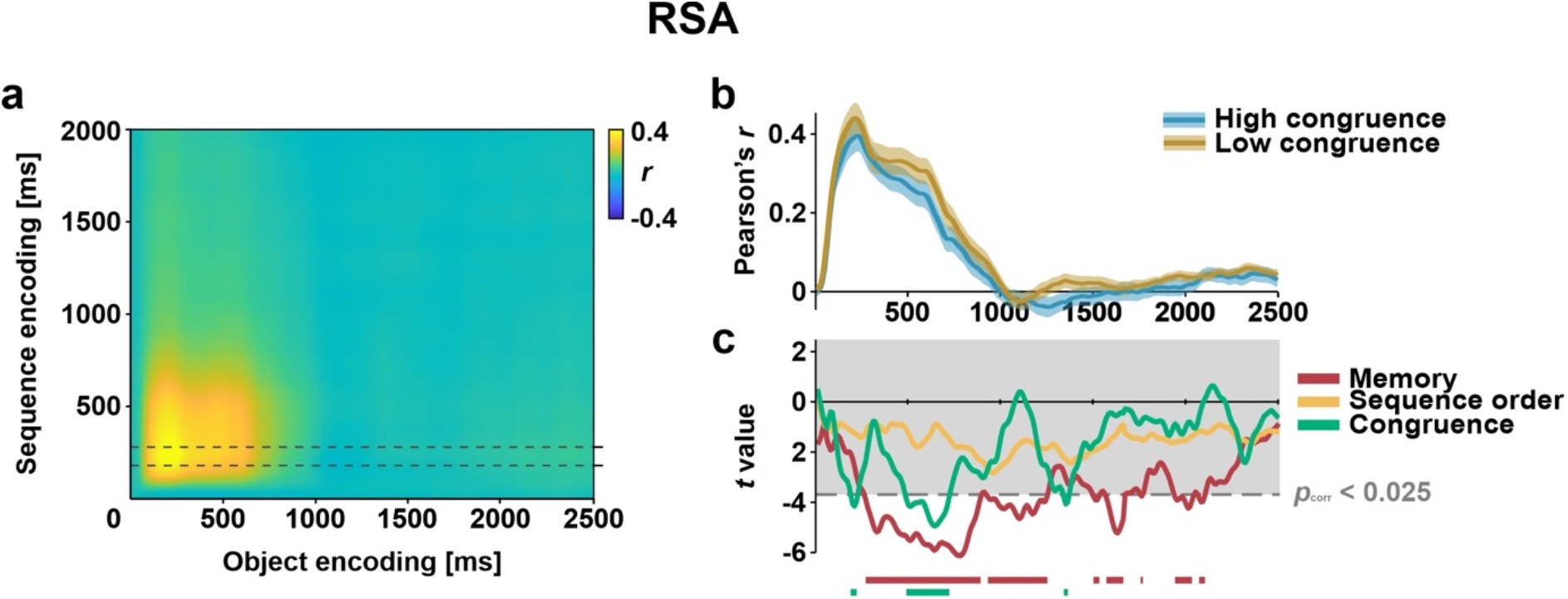
Neural similarity between episode and object. **(a)** Time-resolved degree of neural similarity between EEG patterns elicited by picture images from the episodic sequence and EEG patterns elicited during the encoding of the associated picture object. Grey dashed line shows the ± 50 ms time window where similarity reached the peak (190ms – 290ms at encoding time). **(b)** Neural similarity values for the High- and Low-congruence conditions averaged across participants. Shaded area represents SEM of the participants’ sample. **(c)** *t*-statistics from the output of LMM on similarity value along picture-object encoding period. Grey dashed line marks the one-sided significant threshold after Bonferroni correction. Grey shade shows the area with *p*_corrected_ > 0.05 (two-tailed).

To assess whether the degree of neural reactivation observed during the encoding of picture objects was modulated by their perceived congruency with the encoding episodic context and their later memory accessibility at test, we ran an LMM analysis including neural similarity, participant congruence rating and recognition accuracy at test. We also included a variable accounting for the picture order of pictures in the episodic sequence to control for the possibility that any effect could be accounted for by the temporal proximity of the encoding pictures of the preceding episode with the picture object. To do so, we first identified on the similarity matrix the exact time point where the similarity value reached the peak across participants (i.e., at 240 ms from episodic sequence picture onset and at 220 ms from object picture onset). We then ran a single trial LMM analysis on the averaged similarity value around ± 50ms around the peak (**Figure 5b**). We found that the degree of neural reactivation correlated negatively with the participants’ ability to correctly recognise an item at test. The predictive negative relationship between neural reactivation and memory started to be significant at 270 ms from picture-object onset and remained significant throughout almost the entire epoch until 2100 ms (**Figure 5c**). In addition, we also found a negative correlation between neural similarity and congruence rating. However, the significant effects were more transient but comparably distributed along with the object-picture encoding epoch. Notably, the first significant timepoint was very early, at 190 ms from picture onset, which preceded the relationship effects between neural similarity and memory (**Figure 5c**). A more persistent negative correlation was also found later, between 490 ms – 720 ms and at 1340 ms from picture onset. Finally, no significant point exceeded the statistical significance threshold for the variable picture order position, indicating that the significant relationship between neural similarity and memory and congruence was not driven by specific neural similarity measures between picture objects and pictures of the episode.

## Discussion

In this study, we tested whether the perceived incongruence of an item from an unfolding event and its impact on memory relied on the disruption of neural stability patterns preceded by the neural reactivation of the memory representations of the just-encoded event. Our findings, derived from combining behavioural data from 3 different experiments and the implementation of multivariate pattern analysis on EEG signal during encoding of one of them, confirmed our hypothesis by showing that neural stability patterns gradually increase throughout the ongoing exposure to a schema-consistent episodic narrative, and that the brain stability pattern is interrupted when the encoding of an object of the event is lowly congruent within the ongoing schema representation. We found that the decrease in neural stability for low-congruence items was best seen at ∼1000 ms from object-encoding onset when compared to high-congruence items and that this effect was preceded by an enhanced N400 ERP and an increased degree of neural reactivation of the just-encoded episode for low-congruence items observed between ∼200 to 1000 ms from picture onset. Current results offered new insights into the neural mechanisms and their temporal orchestration that are rapidly engaged during online encoding of schema-consistent episodic narratives and the detection of incongruencies.

Central in our findings is that the degree of neural reactivation of the encoded episode by the final target object correlated negatively with the perceived congruence and the participant’s ability to later recognise the picture in a memory test. The notion that memory reactivation benefits memory formation is well established in previous research. Most of it showed that the reactivation strength drives long-term memory formation by mimicking neural replay phenomena thought to promote rapid consolidation processes seen in rodent studies (i.e., Carr et al., 2011). Other studies have revealed that when novel encoding inputs reactivate previously encoded information that overlaps in content, the long-term memory representations of the two event contents become integrated, promoting generalisation and adaptive behaviour (Shohamy and Wagner, 2008; Schlichting and Preston, 2015). Despite the notion that memory reactivation may potentially benefit memory formation, another set of findings described the opposite effect. These studies found that memory reactivation of overlapping memories may yield competition between the two, resulting in interference with a negative impact on memory (Kuhl et al., 2011). In line with this view, we found a negative correlation between object-picture memory accuracy and the degree of elicited memory reactivation of the preceding event, suggesting an interference between the encoding of an incongruent item and the simultaneous memory reinstatement of the just encoded event.

An additional possible explanation for the proactive interference effect found between memory reactivation and incongruence detection is that surprise itself creates an event boundary (Antony et al., 2021), sectioning off the preceding and the current elements as distinct events in memory. In fact, theoretical models propose that mnemonic prediction errors would promote the encoding of distinct memory traces (McClelland et al., 1995; Love et al., 2004; Gershman et al., 2014; Frank et al., 2020). That is, events that violate our expectations should be allocated a unique memory representation distinct from other existing memories. This may facilitate memory for the unexpected event while mitigating interference with existing memories that may still be relevant. Our findings that high- and low-congruence items were retrieved with similar accuracy when memory was tested with a detailed visual representation in Experiment 2 and rejecting lure items in Experiment 1 would fit this idea, and it would also converge with recent behavioural findings that mnemonic prediction errors do not increase gist-based mistakes of identifying similar lures as old (Bein et al., 2021).

Our findings that surprising or unexpected elements of the unfolding experience elicited the rapid reinstatement of the just-encoded picture sequence are in line with previous findings that showed that sudden shifts in an ongoing episodic context (i.e., event boundaries) induce the rapid reactivation of preceding episodic information (Sols et al., 2017; Silva et al., 2019). Event boundaries are thought to represent moments in time whereby a continuous stream of incoming information is segmented into different memory units (Zack et al., 2011). In this model, the process of event segmentation starts when our current understanding of the world is destabilised by a new observation that does not fit our current expectations. Viewed from this perspective, high levels of surprise (defined here as a high degree of inconsistency of the object picture to the schema or internal model representation activated during the preceding episodic sequence) refer to a substantial change in our understanding of the current inputs from experience. This engenders additional resources to re-evaluate the current internal model in the face of the new observation, which may benefit from the greater reactivation of the memory representations to resolve it.

In summary, the current study offers three new findings. It shows that the detection of low-congruence elements of an episodic experience elicited a rapid memory reactivation of the just-encoded event episodic information; that this is concomitant to a mnemonic prediction error signal during encoding; and that the result of this computation leverages the disruption of stable patterns of neural activity elicited during the schema-consistent episodic event. These findings inform us about the rapid but sequential structure of the distinct neural mechanisms supporting the detection of incongruencies during encoding and their consequences on memory. We speculate that these same processes may take place in realistic scenarios of our everyday experience.

## Acknowledgments

We thank Luz Bavassi for their helpful discussions on earlier versions of the manuscript and Dàlia Dohler for their drawings in Figures 1 and 3. This work was supported by the Spanish Ministerio de Ciencia, Innovación y Universidades, which is part of Agencia Estatal de Investigación (AEI), through the project PID2019-111199GB-I00 (Co-funded by European Regional Development Fund. ERDF, a way to build Europe), to L.F. We thank CERCA Programme/Generalitat de Catalunya for institutional support.

